# Involvement of epithelial-mesenchymal transition genes in small cell lung cancer phenotypic plasticity

**DOI:** 10.1101/2022.09.09.507376

**Authors:** Sarah M. Groves, Nicholas Panchy, Darren R. Tyson, Leonard A. Harris, Vito Quaranta, Tian Hong

**Affiliations:** Department of Biochemistry, Vanderbilt University, Nashville, TN 37235, USA; Department of Biochemistry & Cellular and Molecular Biology, The University of Tennessee, Knoxville, Knoxville TN 37996, USA; Department of Pharmacology, Vanderbilt University, Nashville, TN 37235, USA; Department of Biomedical Engineering, University of Arkansas, Fayetteville, AR 72701, USA; Interdisciplinary Graduate Program in Cell and Molecular Biology, University of Arkansas, Fayetteville, AR 72701, USA; Cancer Biology Program, Winthrop P. Rockefeller Cancer Institute, University of Arkansas for Medical Sciences, Little Rock, AR 72205, USA; National Institute for Mathematical and Biological Synthesis, Knoxville, TN 37996, USA

## Abstract

Small cell lung cancer (SCLC) is an aggressive cancer recalcitrant to treatment, arising predominantly from epithelial pulmonary neuroendocrine (NE) cells. Intra-tumor heterogeneity plays critical roles in SCLC disease progression, metastasis and treatment resistance. At least five transcriptional SCLC NE and non-NE cell subtypes were recently defined by gene expression signatures. Transition from NE to non-NE cell states and cooperation between subtypes within a tumor likely contribute to SCLC progression by mechanisms of adaptation to perturbations. Therefore, gene regulatory programs distinguishing SCLC subtypes or promoting transitions are of great interest. Here, we systematically analyze the relationship between SCLC NE/non-NE transition and epithelial to mesenchymal transition (EMT)—a well-studied cellular process contributing to cancer invasiveness and resistance—using multiple transcriptome datasets from SCLC mouse tumor models, human cancer cell lines and tumor samples. The NE SCLC-A2 subtype maps to the epithelial state. In contrast, SCLC-A and SCLC-N (NE) map to a mesenchymal state (M1) that is distinct from the non-NE mesenchymal state (M2). The correspondence between SCLC subtypes and the EMT program paves the way for further work to understand gene regulatory mechanisms of SCLC tumor plasticity with applicability to other cancer types.

## Introduction

Small cell lung cancer (SCLC) is one of the most aggressive human cancers, characterized by early metastasis and acquisition of therapeutic resistance (Gazdar et al., 2017). SCLC tumors are neuroendocrine (NE) tumors that arise in the lung epithelium, and pulmonary NE cells (PNEC) are considered their normal counterparts (Borges et al., 1997; Song et al., 2012). However, multiple reports have demonstrated considerable intratumoral heterogeneity in SCLC (Calbo et al., 2011; Gay et al., 2021; Lim et al., 2017; Shue et al., 2018; Stewart et al., 2020; Wooten et al., 2019). There is accumulating evidence that SCLC NE cells can transition into non-NE states (Ireland et al., 2020; Lim *et al*., 2017; Stewart *et al*., 2020). Furthermore, prior studies suggest cooperative interactions occur between NE and non-NE cells during SCLC tumor progression (Calbo *et al*., 2011; Ko et al., 2021; Kwon et al., 2015). An open question is whether transitions from NE to non-NE subtypes, and possibly vice versa, are driven by intrinsic factors (e.g., transcription factor network dynamics (Wooten *et al*., 2019)), extrinsic factors (i.e., microenvironmental influences, such as hypoxia or therapeutic agents) (Gay *et al*., 2021; Udyavar et al., 2017), or a combination of both.

A standard to classify SCLC cells based on expression levels of four key transcription factors (TFs), *ASCL1, NEUROD1, YAP1*, and *POU2F3*, was recently adopted (Rudin et al., 2019; Yazawa, 2015). Based on the most abundantly expressed TF, or an underlying signature, three NE (A, A2 and N) and two non-NE (Y and P) subtypes have been proposed (Groves et al., 2022; Wooten *et al*., 2019). An additional subtype enriched in inflammatory genes, SCLC-I, has also been described (Gay *et al*., 2021). However, the gene regulatory mechanisms that generate and maintain the molecular and phenotypic identities of SCLC subtypes remain unclear. Of particular interest are mechanisms underlying NE to non-NE phenotypic transitions since they may directly contribute to tumor progression (Ireland *et al*., 2020; Lim *et al*., 2017; Wu et al., 2021; Zhang et al., 2018). Because PNECs and SCLC cells are of epithelial origin (endodermally derived) (Noguchi et al., 2020) and SCLC tumors are highly metastatic, we hypothesize that similarities may exist between NE/non-NE transitions and the well-studied epithelial-mesenchymal transition (EMT). EMT is a cellular process in which epithelial cells lose tight cell-junction and gain the ability to migrate, characteristic features of metastatic spread (Kalluri and Weinberg, 2009). EMT also contributes to tumor drug resistance (Fischer et al., 2015; Krebs et al., 2017; Mani et al., 2008), and EMT transition has been associated with drug resistance in human SCLC tumors (Stewart *et al*., 2020; Sutherland et al., 2022). Theoretical and experimental studies have revealed that EMT is a multi-stage process and the transcriptional programs (e.g., subsets of mesenchymal signature genes) activated in different contexts can be diverse (Cook and Vanderhyden, 2020; Hong et al., 2015; Lu et al., 2013; Watanabe et al., 2019; Ye et al., 2015; Zhang et al., 2014). For example, different EMT stimulants can activate diverse transcriptional programs (Cook and Vanderhyden, 2020). Thus, while multiple reports have suggested a role for EMT in SCLC phenotypic plasticity, drug resistance, and metastasis, to the best of our knowledge, a systematic analysis of the relationships between NE and non-NE SCLC subtypes and EMT has not been previously reported.

In this paper, we use accepted scoring metrics for epithelial (E) and mesenchymal (M) programs to position NE and non-NE SCLC subtypes within the EMT spectrum. We find that the A2 subtype is strongly epithelial-like (scores high along the E axis) and weakly mesenchymal-like (scores low along the M axis), supporting it as the most E-like subtype of SCLC. In contrast, the non-NE subtypes (P and Y) score higher on the M axis, while being lowest along the E axis. Surprisingly, the NE subtypes A and N consistently score significantly lower than A2 along the E axis. Moreover, transcripts of some mesenchymal genes, such as ZEB1, are more abundant in N than in non-NE subtypes, suggesting the existence of distinct mesenchymal gene programs across SCLC subtypes. These findings support the involvement of EMT programs in SCLC phenotypic heterogeneity and provide potential mechanistic insights into NE/non-NE transitions.

## Results

### The SCLC-A2 subtype is highly enriched in epithelial gene expression

We first analyzed a time course of single-cell RNA-sequencing (scRNA-seq) dataset from an SCLC genetically engineered mouse model (GEMM) with an overactive Myc oncogene (*Rb1^fl/fl^; Trp53^fl/fl^*; Lox-Stop-Lox [LSL]-*Myc^T58A^*, RPM) (Ireland *et al*., 2020). The 15,138 single cells were annotated with the SCLC subtypes using gene expression signatures and archetype analysis, as recently described (Groves *et al*., 2022). An SCLC signature comprising 105 genes that distinguishes the five SCLC subtypes was used to classify individual cells relative to extreme expression patterns of NE and non-NE subtypes, with some subpopulations enriched in multiple subtype signatures, resulting in four classes (A/N, A2, P/Y, and Y) (see Methods) (Groves *et al*., 2022). In parallel, using normalized gene expression, we computed the epithelial (E) and mesenchymal (M) scores for this dataset. The scores were computed using single-sample gene set enrichment analysis (ssGSEA) with a list of 232 epithelial (E) associated genes and 193 mesenchymal (M) associated genes (see Methods) (Deshmukh et al., 2021; Krug et al., 2019; Tan et al., 2014). In the EMT spectrum depicted by the E and M scores, we observed a striking difference between the position of the NE A2 subtype and other NE and non-NE subtypes (Figure 1A). The A2 cells had a significantly higher E score and a significantly lower M score compared to other subtypes (*p*<10^−5^ for each comparison). Consistent with the E/M scores, we found that A2 cells had significantly higher levels of *Cdh1* (coding E-cadherin) (*p*<10^−5^), a widely used E marker gene crucial for cell adhesion, compared to other cells (Figure 1A inset). These results indicate that gene expression in the A2 subtype is most restricted to an epithelial phenotype, whereas, in all other SCLC subtypes (NE and non-NE), M gene expression appears to be permitted to various extents.

**Figure 1.**
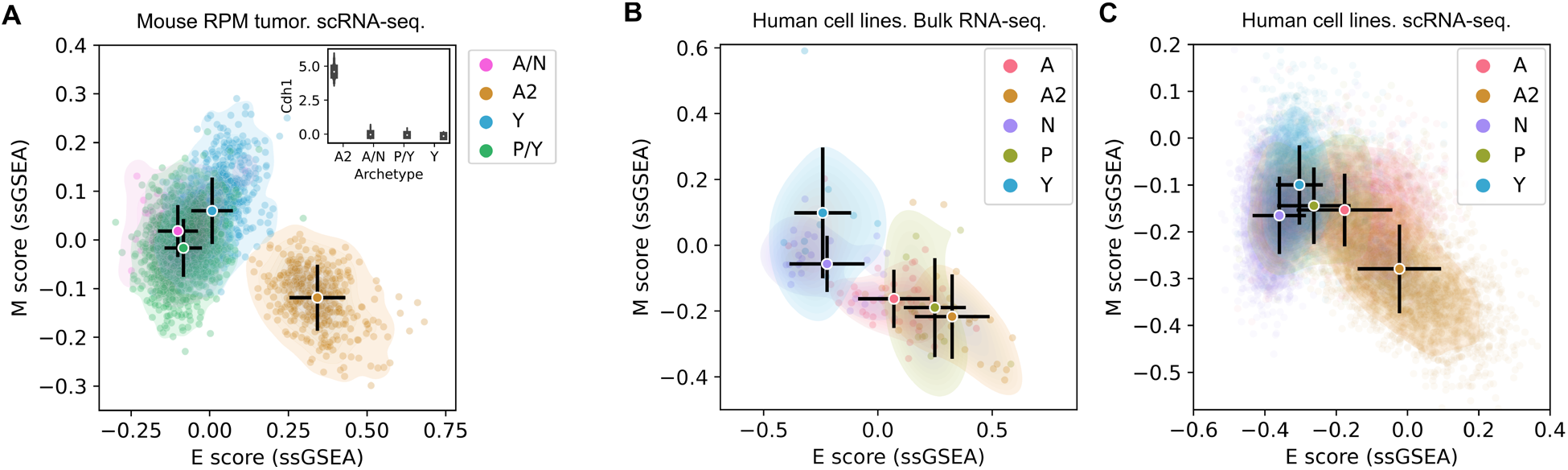
Distributions of SCLC cells across the epithelial and mesenchymal spectrum. E and M scores of **A**. 15,138 mouse *Myc*-overexpressing SCLC GEMM tumor cells in an *ex vivo* model; **B**. 120 SCLC cell lines; and **C**. 13,945 cells from eight SCLC cell lines. Scores were computed with ssGSEA and 425 previously identified EMT genes. Circles and black bars indicate means and standard deviations. Inset in A shows normalized expression of Cdh1 in four classes of cells.

We next asked whether the association between the A2 subtype and the epithelial lineage in RPM mouse SCLC tumor cells extends to human SCLC cells. We first analyzed bulk RNA-seq data from 120 human SCLC cell lines assigned with subtype identities (see Methods). Several A2 cell lines (e.g., DMS53) were located in the extreme E position on the EMT spectrum (Figure 1B), in agreement with A2 cells from RPM tumors (Figure 1A). As a group, A2 cell lines had the highest mean E score and the lowest M score among SCLC subtypes, although these scores were closer to A and Y cells than in the RPM tumor cell data (Figure 1B). This partial overlap could be due to cell-cell heterogeneity within cell lines that is not resolved in bulk RNA-seq data. Therefore, we visualized 13,945 single-cell transcriptomes from eight human SCLC cell lines (Groves *et al*., 2022) with the same E/M projection method. In this dataset, A2 cells are more distinct from other subtypes than in the bulk RNA-seq data, and the separation is more similar to the RPM tumor cell data (Figure 1C). Interestingly, in both bulk and single-cell RNA-seq datasets for SCLC cell lines, the NE subtype A has an intermediate EMT expression profile (Figure 1B, C).

The strong association between SCLC subtypes and EMT progression raises the question of whether this connection is trivially due to a sharing of genes between the 105 SCLC subtype signature gene set and the 425 EMT signature gene set. However, there are only 20 genes appearing in both gene sets, indicating limited overlap (Jaccard index 0.039). Furthermore, excluding the 20 overlapping SCLC signature genes from the EMT gene set had minimal effect on the distributions of SCLC cells in the EMT spectrum in each of the three datasets (Figure S1). Taken together, our analyses indicate a strong and nontrivial association between SCLC subtypes and EMT progression. In particular, a specific SCLC NE subtype, A2, exhibits phenotypes associated with highly restricted epithelial lineages in mammals.

### Mesenchymal scoring is diverse across non-A2 subtypes

The ssGSEA-based E/M scoring did not show any dramatic difference among A, N, P, and Y (i.e. non-A2) subtypes (Figure 1). However, the expression of individual M genes, such as *Vim*, differed significantly among non-A2 subtypes (Figure 2A, B). This suggests the possibility that M genes have divergent expression patterns across the SCLC subtypes which may mask the ssGSEA-based scoring. Recent studies in tumors and cell lines also suggested diversity of EMT programs (Cook and Vanderhyden, 2020; Pastushenko et al., 2018; Watanabe *et al*., 2019). For example, activation of a subset of M genes depends on EMT transcription factor ZEB1, while that of other M genes are activated via ZEB1-independent pathways (Watanabe *et al*., 2019). To determine potentially distinct M programs in SCLC cells, we used an alternative scoring method based on nonnegative principal component analysis (nnPCA), which generates leading PCs that describe the variance for each gene set (see Methods). With the single-cell data from both the mouse RPM model and the SCLC cell lines, we noticed that the first PC for M scores explained less variance compared to the first PC for E scores (Figure 2C, D top). Furthermore, the first two M PCs produced comparable variances explained. These results indicate that M gene expression is spread evenly over at least two orthogonal dimensions, suggesting diversity of M scoring and warranting further investigation. Interestingly, M scores obtained from the first and second PCs ranked the SCLC subtypes differently: A/N subtypes have *higher* scores than Y subtype cells based on the first M PC (nnPC1) but *lower* scores than Y subtype with the second M PC (nnPC2) (Figure 2C, D scatter plots). In fact, two representative M genes—*Vim* (the gene encoding vimentin, an intermediate filament protein component of mesenchymal cell cytoskeletons) and *Zeb1* (a widely studied EMT transcription factor)—had an anticorrelated pattern between A/N and Y subtypes in the RPM dataset (compare Figure 2E to Figure 2A) (Mendez et al., 2010; Wellner et al., 2009). Although this anticorrelation was less prominent in the cell line data (compare Figure 2F to Figure 2B), distinct *Vim* and *Zeb1* expressions of SCLC subtypes were observed.

**Figure 2.**
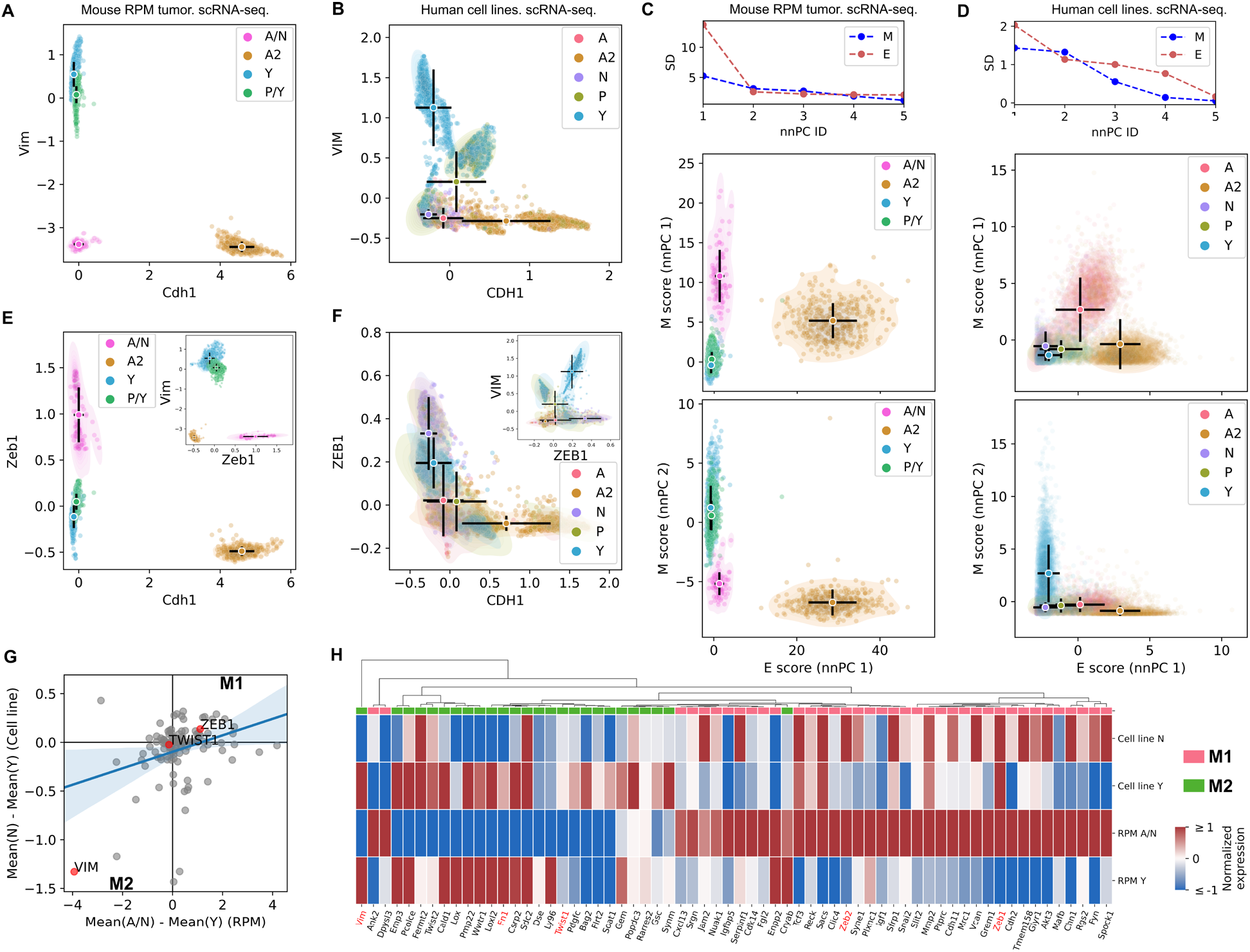
Divergence of mesenchymal gene expressions in SCLC subtypes. **A, B**. Normalized expression levels of Vim (M gene) and Cdh1 (E gene) in 15138 mouse *Myc*-driven tumor cells (A) and in 13945 cells from eight SCLC cell lines (B). Imputed data were plotted in B for visual aid. Circles and black bars show means and standard deviations of SCLC subtypes. **C, D**. Line plots: nnPCA was performed. Plots show standard deviations of SCLC subtype means for mouse and human SCLC cells from the top 5 PCs. Scatter plots: nnPCA-based E and M scores. **E, F**. Normalized expression levels of Zeb1 (M gene) and Cdh1 (E gene) in 15138 mouse *Myc*-driven tumor cells (E) and in 13945 cells from 8 SCLC cell lines (F). Imputed data were plotted in F for visual aid. **G**. Mean differences between N cells (A/N cells for mouse) and Y cells in individual M gene’s expression in mouse and human SCLC cells. **H**. A heatmap showing the diversity of M gene expression across N (A/N) and Y subtypes.

Overall, like the case of *Vim* and *Zeb1*, the mean differences in expression of M genes between A/N subtypes and Y subtype were diverse, and these differences were positively correlated between the RPM data and the cell line data (Figure 2G, Pearson correlation coefficient 0.29, *p*<10^−5^). Some M genes, such as *Zeb1*, had higher expression in the A/N subtypes than in the Y subtype, while some others, such as *Vim*, had the opposite pattern (Figure 2G, H). We defined these two groups of M genes as M1 and M2 respectively (FDR<0.05 for N-Y differences). We performed gene ontology analysis and found that, while 20 enriched biological processes were shared between the two groups, M1 genes were uniquely enriched in more than 200 processes including ‘negative regulation of cell adhesion involved in substrate-bound cell migration’ (fold enrichment >100, FDR=0.005), whereas M2 were uniquely enriched in 11 processes including ‘extracellular matrix organization’ (fold enrichment =14.08, FDR=0.02) (Tables S1 and S2). As these two groups of processes may both contribute to enhanced cell motility, this result suggests that NE and non-NE subtypes may use distinct strategies to achieve mesenchymal-like cellular functions such as cell migration. In summary, our result supports the functional divergence of the M genes differentially expressed in NE and non-NE SCLC subtypes.

### The highly epithelial A2 subtype and diverse M gene expression patterns are detectable in human SCLC tumor

The distinction between the A2 and A subtypes has generally not been considered in previous studies by other investigators. For example, Chan et al. profiled 54,523 individual SCLC cells from 19 human samples and classified them into A, N, and P subtypes (note the absence of the Y subtype) (Chan et al., 2021). We therefore asked whether our analytical framework can be used to discover previously unknown, extremely epithelial-like cells in this human SCLC tumor dataset. We first performed E and M scoring for this dataset (Figure 3A) and found the A cells (as subtyped in Chan et al., 2021) fell into two distinct groups, one of which was highly E-like (Figure 3B, red arrow). We hypothesized that the cells labeled as A subtype by Chan et al. include both extreme E-like (A2) and EMT-intermediate-like (A) subtypes. We re-subtyped the SCLC cells from Chan et al. using our subtype gene signatures (see Methods), and we found a significant fraction of the previously labeled A cells had high scores for the A2 subtype and low scores for the A subtype, which we labeled A2* (Figure 3C, D, S2). In particular, the distinct group of E-like cells corresponds to the group of cells that had high A2 scores and low A scores (Figure 3C, D. Figure S2). Interestingly, the cell cluster with high A2 scores corresponds to a unique tumor sample (Figure 3A, pink) with a unique treatment type (Figure S3). We found that the A2 scores of these 11,056 A2-enriched SCLC cells were strongly positively correlated with their E scores (nnPCA performed with RU1108 cells alone) (Figure 3E) (Spearman correlation coefficient *s_r_* = 0.69). In contrast, A scores were negatively correlated with E scores in these cells (Figure 3E) (*s_r_* = −0.23). In addition, A2 scores had a moderately negative correlation with M scores (nnPCA) (*s_r_* = −0.17) (Figure 3F).

**Figure 3.**
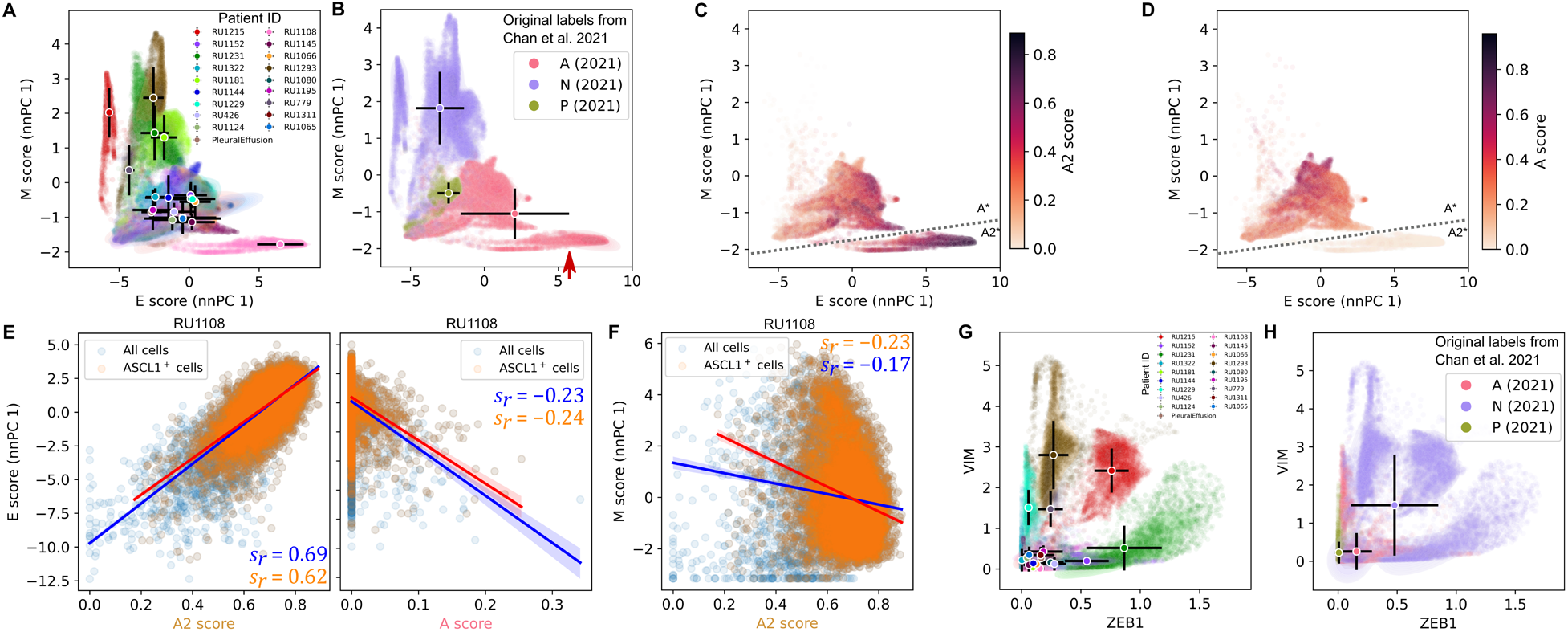
Detection of epithelial- and mesenchymal-like SCLC subtypes in human tumor cells. **A**. Scatter plot shows nnPCA-based E and M scores for 54,523 SCLC cells (Chan *et al*., 2021). Color code represents each of the 19 patients. Circles and black bars show means and standard deviations of cell scores for each patient. **B.** Same data as in A with SCLC subtype labels obtained from Chan et al. (2021). Circles and black bars show means and standard deviations of scores across the previously determined SCLC subtypes. **C**, **D**. Scatter plots show A2 and A subtype scores in the previously determined SCLC-A subtype cells from the data in B. **E, F**. Transcriptome data of 11,056 SCLC cells from tumor RU1108 of the Chan et al. study (Chan *et al*., 2021) were projected onto indicated score axes. Linear regression lines and confidence intervals were obtained for all cells and ASCL1^+^ cells. *s_r_* is Spearman correlation coefficient. **G**. Normalized expression of *ZEB1* and *VIM* in the same dataset as in A. Color code represents each of the 19 patients as in A. **H**. The same data as in G but color labels are defined by SCLC subtype shown in B.

Systematic analysis for divergent M programs could not be performed due to the absence of any Y subtype cells in the dataset. Nonetheless, heterogeneous expressions of M genes *ZEB1* and *VIM* were observed within the N subtype (Figure 3G). Furthermore, three M-like cell clusters with similar E and M score rankings (Figure 3A, red, dark green, and brown) had dramatically different profiles of *VIM* and *ZEB1* expression (Figure 3G) even though all were classified as N (Figure 3H). These results further support the association between SCLC and EMT programs and reveal a previously underappreciated connection between SCLC tumor cell heterogeneity and EMT.

### Intratumor heterogeneity of SCLC indicates strong A2-epithelial association at single-cell level

The A2-epithelial (A2-E) association in human tumors described in the previous section relies on gene set analyses across multiple tumors, where all cells within each tumor were classified into a single subtype (Chan *et al*., 2021) (Figure 3A-D). We next analyzed a scRNA-seq dataset of two human SCLC circulating tumor cell-derived xenografts (CDXs) (Gay *et al*., 2021), which each underwent an EMT-like cell state transition upon cisplatin treatment such that multiple subtypes exist within individual tumors. We first performed subtyping analysis and computed scores of five SCLC subtypes for 5,268 cells (untreated and cisplatin-treated) of tumor SC53. We visualized the SCLC scores in the E-M space obtained from nnPCA (Figure 4A) and found that while there was a distinct, small population corresponding to an M-like state as previously reported (Gay *et al*., 2021), most tumor cells were located in a continuous region with relatively high A2 and A scores (Figure 4A), consistent with their positivity for ASCL1 previously shown. Furthermore, N and P subtypes seem to be absent from this tumor (Figure 4A). Interestingly, although both A and A2 scores were positively correlated with E scores for all cells in the tumor (*s_r_* = 0.43 for A2; *s_r_* = 0.09 for A), only A2 scores had a strong correlation with E scores for cells that express ASCL1 (Figure 4E and F) (*s_r_* = 0.35 for A2; *s_r_* = −0.08 for A). In addition, neither the SC53 tumor cells nor its ASCL1^+^ subpopulation showed a strong correlation between A2 scores and M scores (Figure 4G).

**Figure 4.**
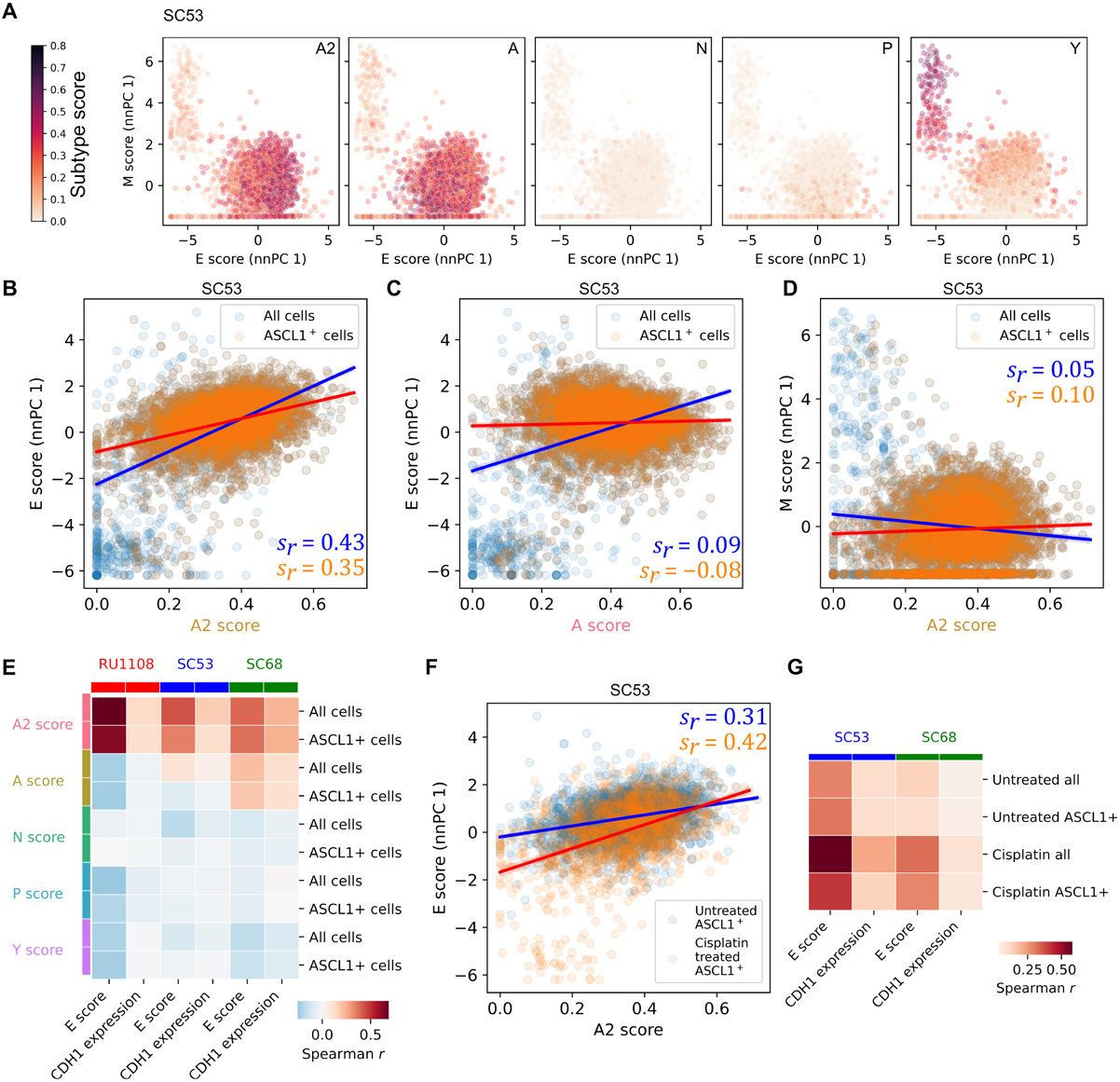
Correlations between A2 and epithelial transcriptional programs in individual human tumor cells. **D.** 5,268 single-cell transcriptomes from SCLC tumor SC53 (Gay *et al*., 2021) were projected onto the E and M score axes using nnPCA. Colors indicate SCLC subtype scores (see Methods). **E-G**. Linear regression lines and confidence intervals were obtained for all cells and ASCL1^+^ cells from data in D. **H**. Datasets of three tumors (top labels) were used to compute the Spearman correlation coefficients between SCLC subtype scores (left labels) and E scores from nnPCA or CDH1 expression levels. All cells and ASCL1^+^ cells were analyzed separately. **I**. Correlations between A2 scores and E scores (nnPCA) in untreated and cisplatin-treated cells of SC53 tumor. **J**. Datasets of SC53 and SC68 tumors were used to compute the Spearman correlation coefficients between SCLC subtype scores (left labels) and E scores from nnPCA or CDH1 expression levels. All cells and ASCL1^+^ cells were analyzed separately. Untreated and cisplatin-treated cells were analyzed separately.

We next extended our analysis to SC68, another tumor that underwent an EMT-like transition upon cisplatin treatment (Gay *et al*., 2021). Similar to SC53, there was a strong positive correlation between A2 and E scores both in all cells in the tumor and in ASCL1^+^ cells, whereas the correlations between other subtypes’ scores and E scores were either significantly weaker or negative (Figure 4H). We also found that the strong A2-E association was partly explained by a positive correlation between A2 scores and the expression of CDH1 (Figure 4H). Again, the A2-E association was not due to shared genes between the two gene sets that we used for scoring (Figure S3). Together, our results show that the intratumor association between A2 and E transcriptional programs is a consensus pattern across RU1108, SC53, and SC68 tumors.

We next asked whether the A2-E correlation in SC53 was driven by cisplatin treatment or intrinsic cell-to-cell heterogeneity. We found that cisplatin-treated ASCL1^+^ cells showed both greater variance in the E-M space and stronger correlation between A2 and E scores compared to untreated cells (Figure 4I) (*s_r_* = 0.31 for untreated cells; *s_r_* = 0.42 for treated cells). Nonetheless, even in the untreated cells, a positive correlation between A2 and E scores was observed (Figure 4I). Therefore, this A2-E association was found across all cells of both tumors before and after treatment (Figure 4J). These results excluded the possibility that the observed A2-E association was due to Simpson’s paradox in which the treatment condition may be a hidden variable. Instead, they suggest that both cisplatin treatment and intrinsic cell-to-cell heterogeneity contribute to the A2-E correlation and that treatment can induce a cell state transition towards an M-like state without turning off ASCL1 transcription.

## Discussion

Understanding the molecular basis for tumor heterogeneity is crucial for developing next-generation therapeutic strategies (Altschuler and Wu, 2010; Jordan et al., 2016; Sáez-Ayala et al., 2013; Shaffer et al., 2017). Recent advances in defining subtypes of SCLC tumors provides such potential (Gay *et al*., 2021), but cellular and molecular mechanisms contributing to phenotypic heterogeneity are still unclear. Interestingly, SCLC tumors have both NE and epithelial characteristics (Stovold et al., 2013). We therefore asked whether SCLC cells hijack EMT, canonically defined as a developmental program (Thiery et al., 2009), to increase invasiveness and drug resistance which has been shown to occur in other cancer types (Fischer *et al*., 2015; Mani *et al*., 2008).

While it was shown that some genes promoting EMT are activated in NE/non-NE transitions of SCLC (Groves *et al*., 2022; Ireland *et al*., 2020; Ito et al., 2017), connections between SCLC subtypes to the EMT spectrum have not been studied in depth. It is also unclear whether SCLC involves multiple (unique) partial EMT states. Through analysis of transcriptome datasets from multiple sources including a mouse tumor model, human cell lines, and human tumor samples, our work reveals a strong correspondence between the EMT spectrum, containing an epithelial state and several divergent mesenchymal states, and recently defined SCLC subtypes. Our study indicates the partial EMT status of the NE subtypes A and N, whereas the NE subtype A2 is fully epithelial-like. In addition, we show divergence of mesenchymal gene expression in N (an NE) and Y (a non-NE) subtypes.

Our analysis shows diverse expressions of mesenchymal factors across SCLC subtypes. This is consistent with recent observations showing EMT is a context-specific dynamic process (Cook and Vanderhyden, 2020). Interestingly, the EMT transcription factor ZEB1 and EMT effector gene VIM have an anti-correlated expression pattern between N and Y subtypes. Although the contribution of ZEB1 to EMT was demonstrated *in vivo* and *in vitro* (Celià-Terrassa et al., 2018; Cieply et al., 2013; Han et al., 2022; Watanabe *et al*., 2019), its high expression may not be required for some partial EMT states. Our previous work showed that expressions of some mesenchymal genes do not require ZEB1 activation, and that the high expression of a group of EMT genes positively controlled by ZEB1, but not TGF-β, is correlated with better prognosis of breast cancer patients (Watanabe *et al*., 2019). It is possible that ZEB1 is transiently required for maintaining a group of NE cells during SCLC progression, and the transitions to more drugresistant subtypes may require the down-regulation of ZEB1.

This work builds on our prior studies that suggested the partial EMT status of N, based on mixed morphological features and expression of *ZEB1, SNAI1*, and *TWIST1* but not *VIM* (Groves *et al*., 2022). Here, we expand this work by analyzing EMT-related gene signatures in this subtype across multiple datasets. While SCLC-N is canonically defined by expression of NEUROD1, characterizing this subtype as a partial-EMT state may lead to new insights regarding its functional role in SCLC tumor progression and metastasis. Further work is needed to understand how enrichment of a mesenchymal signature (M1) in this subtype is related to this partial-EMT characterization.

The heterogeneity of NE and non-NE SCLC subtypes has been observed previously. However, the factors driving transitions between NE and non-NE subtypes are not well characterized. It has long been known that EMT is reversible in both embryonic and postnatal development, and this reversible process can also be triggered in cancer cell lines using either dynamic extracellular signals (e.g. TGF-β) or forced expression of intracellular factors (e.g. TFs or microRNAs) (Cursons et al., 2018; Grande et al., 2015; Hong *et al*., 2015; Watanabe *et al*., 2019). Our findings suggest that the machinery for reversible EMT may also be responsible for the transitions between NE and non-NE cells during the progression of SCLC. Future work is warranted to determine the temporal sequence of activation among EMT TFs, signaling molecules, and subtype-defining factors for SCLC, such that the intrinsic and extrinsic factors contributing to the NE/non-NE transition can be dissected. Overall, by revealing the relationships between NE/non-NE subtypes and EMT progressions, this study will help guide future work to improve our understanding of SCLC tumor heterogeneity and cell state transitions that may be driven by EMT programs.

## Resource availability

### Lead contact

Further information and requests for resources and reagents should be directed to and will be fulfilled by the Lead Contact, Tian Hong.

### Materials availability

This study did not generate new unique reagents.

### Data and code availability

The code generated during this study is available at GitHub (https://github.com/smgroves/EMT_SCLC_project). No new data was generated for this study.

## Methods

### RNA-sequencing data

We obtained and batch-corrected bulk RNA sequencing expression data on SCLC cell lines from the Cancer Cell Line Encyclopedia (50) and cBioPortal (70) (Cerami et al., 2012). Preprocessing of this data was described in Groves et al. (2022).

Single-cell RNA sequencing data was downloaded from Gene Expression Omnibus (GEO) at GSE193959 (human cell lines), GSE149180 (RPM mouse tumor time course), and GSE138474 (human CDX tumor samples) (Gay *et al*., 2021; Groves *et al*., 2022; Ireland *et al*., 2020). Preprocessed human tumor data was downloaded from the Human Tumor Atlas Network deposited by Chan et al. (Chan *et al*., 2021) at Synapse ID syn23630203. Human cell line and RPM mouse tumor datasets were preprocessed as described in Groves et al. (Groves *et al*., 2022), including filtering and normalization of total counts using the Python package *Scanpy* (Wolf et al., 2018), log-transformation using the *log1p* function from the *Numpy* package, and scaling using *Scanpy*. Log-normalized expression levels were used for comparisons (Mann-Whitney U test) between subtypes.

Human CDX data was preprocessed as described in Gay et al. (2022). Briefly, cells were filtered as described (Gay *et al*., 2021)) to remove non-tumor cells. SC53 tumors (before and after cisplatin treatment) were concatenated together, and SC68 tumors were concatenated together. For each dataset (SC53 and SC68), *Scanpy* was used: total counts were normalized by cell, then data was log transformed (log1p). Highly variable genes were determined with min_mean = 0.0125, max_mean 5, and min_dsip = 0.8. A PCA, tSNE, and Leiden clustering were then calculated using *Scanpy*.

### Subtyping of SCLC cells

Single cell subtype labeling was done as described in Groves et al. (Groves *et al*., 2022). Briefly, after archetype analysis was applied to the MAGIC-imputed single cell datasets and vertices were identified (Hart et al., 2015; Van Dijk et al., 2018), we labeled the cells closest to each archetype as that subtype if the archetype score was > 0.95 (each cell receives weighted archetype scores that sum to 1). SCLC archetypal signatures were generated from bulk human cell line and tumor transcriptomics data, giving a matrix of 105 genes by 5 archetypes (SCLC-A, SCLC-A2, SCLC-N, SCLC-P, and SCLC-Y). In order to align the single-cell archetypes with our bulk archetype signatures, we consider the scores for each cell described in the STAR Methods section “Bulk gene signature scoring of single cells using archetype signature matrix” from Groves et al. (Groves *et al*., 2022). For each bulk signature *x* and for each single-cell archetype *a*, we ran the following significance test:

1. Find the mean bulk score *x* for a specialist, *m*.
2. Choose a random sample of size *n_a_*, where *n_a_* is the number of specialists, with replacement from the remaining cells (i.e. cells that are not specialists, including generalists and other specialist cells). Find the mean bulk score for this sample. N.B. Because some time points have very few cells, we sample evenly from each time point to ensure adequate representation across the time points.
3. Repeat this random selection 1000 times.
4. Generate a *p*-value, which is equal to the percentage of means from this random distribution above *m*.
5. Using statsmodels.states.multitest, correct *p*-values for multiple tests. We used the Bonferroni-Holm method to control the family-wise error rate. Consider *q* < 0.1 significant.

Therefore, each archetype was labeled with an SCLC subtype if enriched in that subtype’s signature. For this work, we considered only the cells labeled with such a subtype (i.e. specialists). For the RPM tumor cells, six archetypes were originally found in Groves et al., 2022 (A/N, A2, P/Y, two Y groups, and one archetype not enriched in any signatures). Here, we consider those enriched in SCLC subtype signatures, combining the two Y groups resulting in A/N, A2, P/Y and Y.

### Single sample gene set enrichment analysis (ssGSEA)

ssGSEA was performed to compute the enrichment scores for individual cells (for scRNA-seq data) or individual cell lines (for bulk RNA-seq data) (Deshmukh *et al*., 2021; Krug *et al*., 2019). A list of 232 epithelial signature genes, and a list of 193 mesenchymal signature genes were used to compute the E enrichment score (E score) and M enrichment score (M score) respectively (Cursons *et al*., 2018; Panchy et al., 2022; Tan *et al*., 2014). The combined list of 425 EMT genes has 20 genes that also appear in the 105 signature genes used for SCLC subtyping. To ensure that the inferred relationship between EMT scoring and SCLC subtyping was not simply due to the shared list of genes, we excluded the intersection of the two gene sets and performed additional ssGSEA scoring for each dataset. The patterns of the scoring were not altered by the exclusion.

### Non-negative principal component analysis (nnPCA)

To quantify the divergent progression of EMT across the subtypes of SCLC cells, we used a second approach to compute the E and M scores. We performed nnPCA using the gene sets mentioned above (Panchy et al., 2021; Panchy et al., 2022). nnPCA determines the approximately orthogonal axes with non-negative coefficients (loadings) for features (genes). Variances of projections of data points (cell or cell lines) onto these axes are maximized via an optimization method (Sigg and Buhmann, 2008). To select the principal components (PCs) that best represent the EMT programs across SCLC subtypes, we used two criteria to rank the PCs in a semi-supervised manner. We first selected the top five PCs that have the highest variances explained for individual samples (cells or cell lines). Among the five PCs, we re-rank them based on the variances of means of individual SCLC subtypes. Similar to ssGSEA, we excluded the EMT gene set and the SCLC gene set and performed additional nnPCA scoring for each dataset. The ranks of the PCs and the patterns of the nnPCA were not altered by the exclusion.

## Acknowledgement and Funding

This work was supported by funds from National Institutes of Health R01GM140462 (T.H.), R50CA24378 (D.R.T.), U54CA217450 (V.Q and S.M.G), K22CA237857 (L.A.H.), and from NSF DGE-1445197 (S.M.G).

## Supplementary Figures

**Figure S1.**
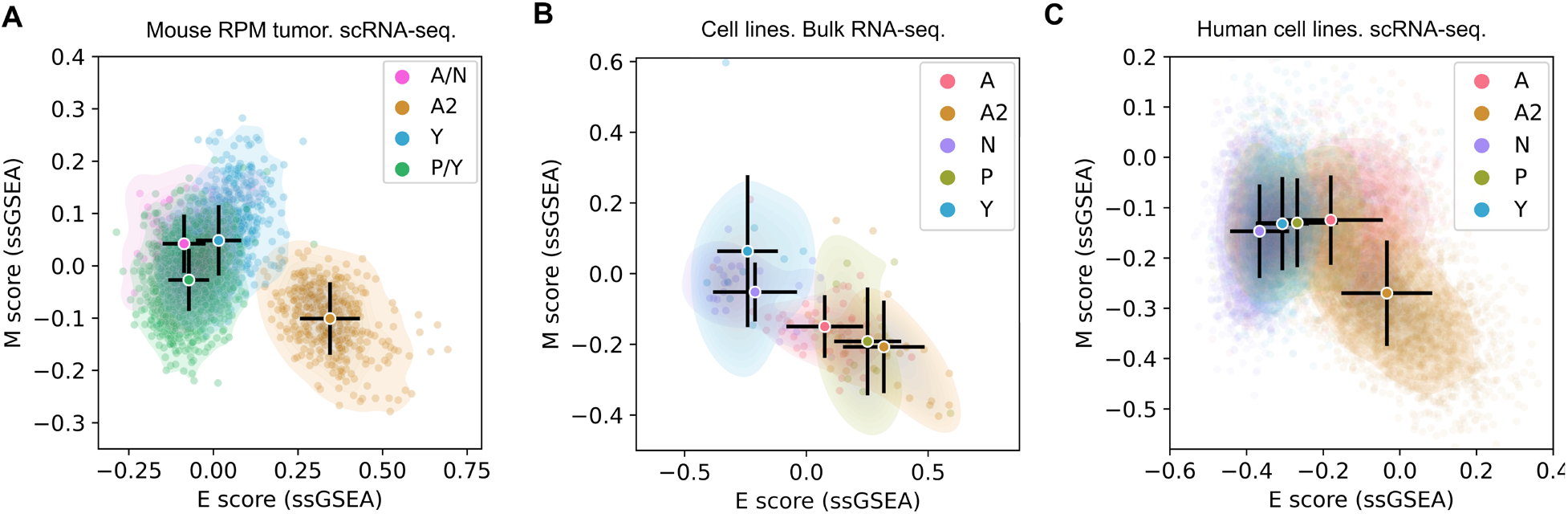
Distributions of SCLC cells across the epithelial and mesenchymal spectrum using non-overlapping gene sets. **A**. E and M scores of 15138 mouse *Myc*-driven tumor cells in an *ex vivo* model. Archetype analysis was performed to determine generalist cells and specialist cells of four types (SCLC subtypes). Scores were computed with ssGSEA and 405 previously identified EMT genes that were not used for SCLC subtyping. **B**. E and M scores of 120 SCLC cell lines computed with the same method as in A. **C**. E and M scores of 13945 cells from 8 SCLC cell lines computed with the same method as in A.

**Figure S2.**
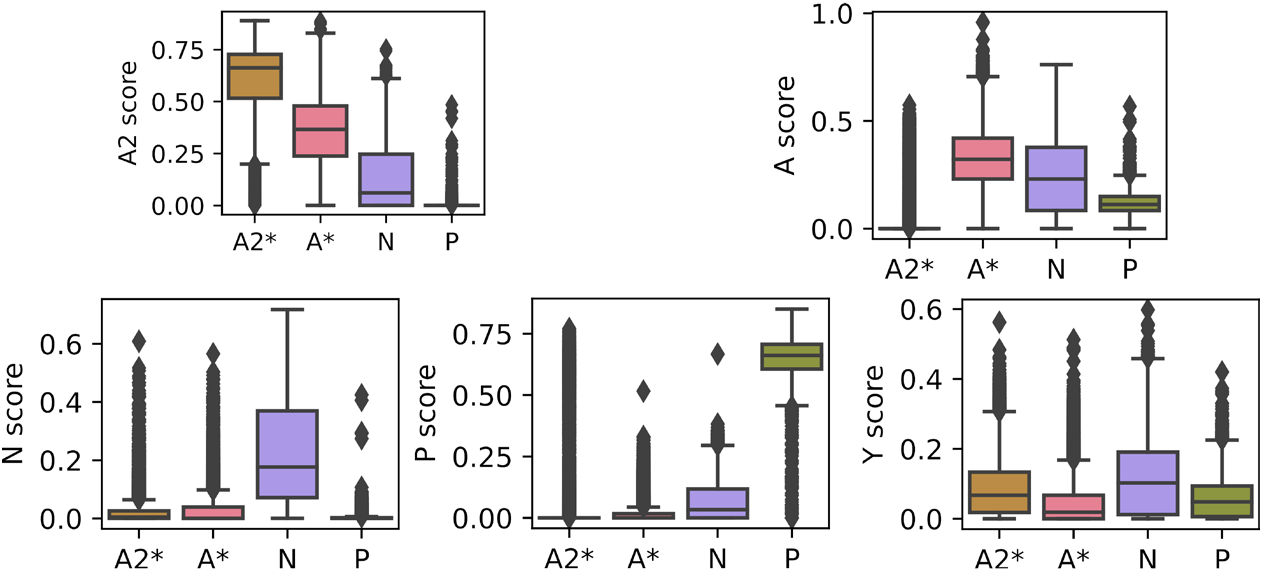
Subtype scores for human tumor cells. Boxplots show subtype scores including newly defined A* and A2* subtypes based on the threshold indicated in Figure 3C and D, as well as N and P subtypes defined by Chan et al. (Chan *et al*., 2021).

**Figure S3.**
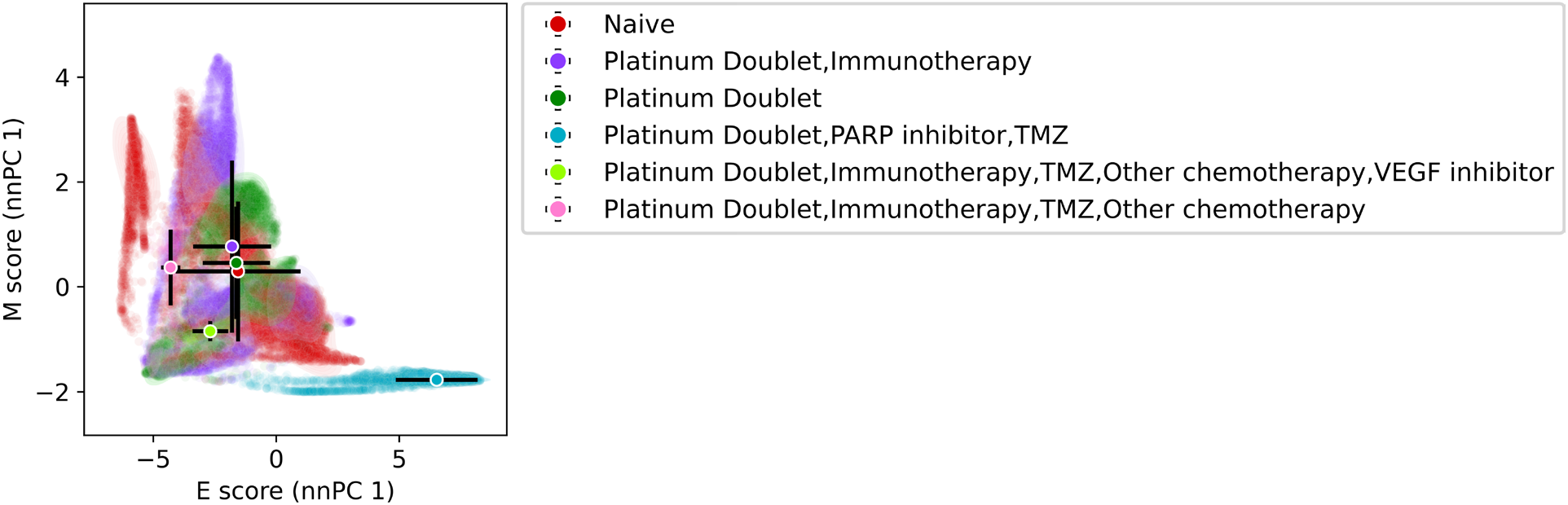
Treatments of human tumor cells visualized in EMT space. Scatter plot shows nnPCA-based E and M scores for 54,523 SCLC cells (Chan *et al*., 2021). Color code represents each of the treatment types. Circles and black bars show means and standard deviations of cell scores for each treatment type.

**Figure S4.**
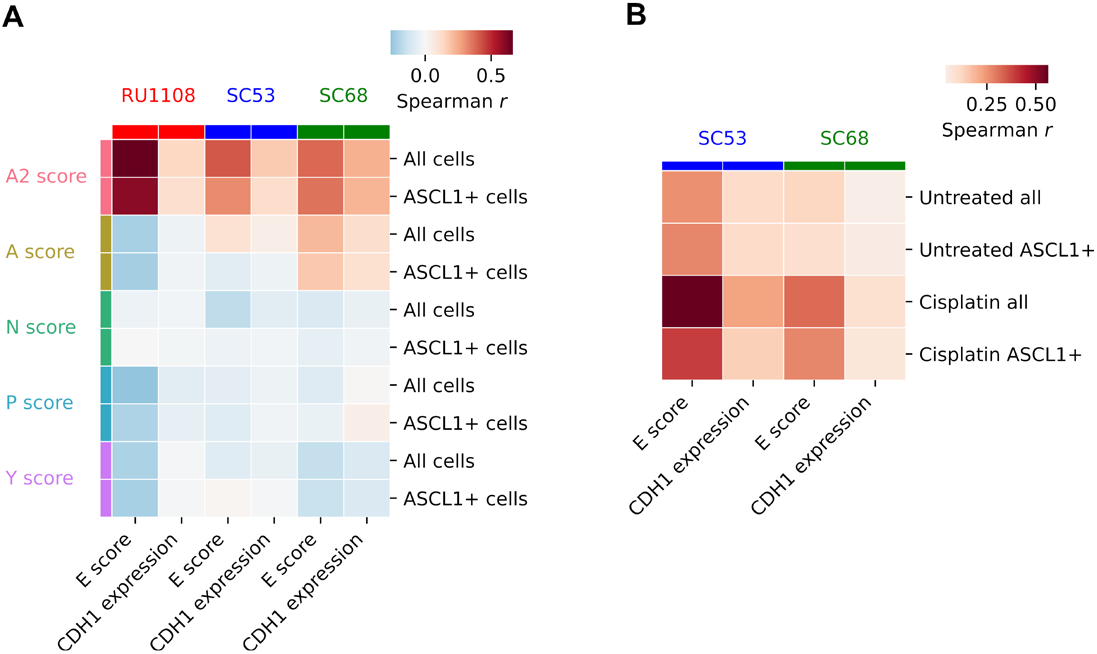
Correlations between A2 and epithelial transcriptional programs in individual human tumor cells using non-overlapping gene sets. **A**. Datasets of three tumors (top labels) were used to compute the Spearman correlation coefficients between SCLC subtype scores (left labels) and E scores from nnPCA or CDH1 expression levels. All cells and ASCL1^+^ cells were analyzed separately. **B**. Datasets of SC53 and SC68 tumors were used to compute the Spearman correlation coefficients between SCLC subtype scores (left labels) and E scores from nnPCA or CDH1 expression levels. All cells and ASCL1^+^ cells were analyzed separately. Untreated and cisplatin-treated cells were analyzed separately.

