## Supplementary figures and images for "Involvement of epithelial-mesenchymal transition genes in small cell lung cancer phenotypic plasticity"

### Figure_S1.png

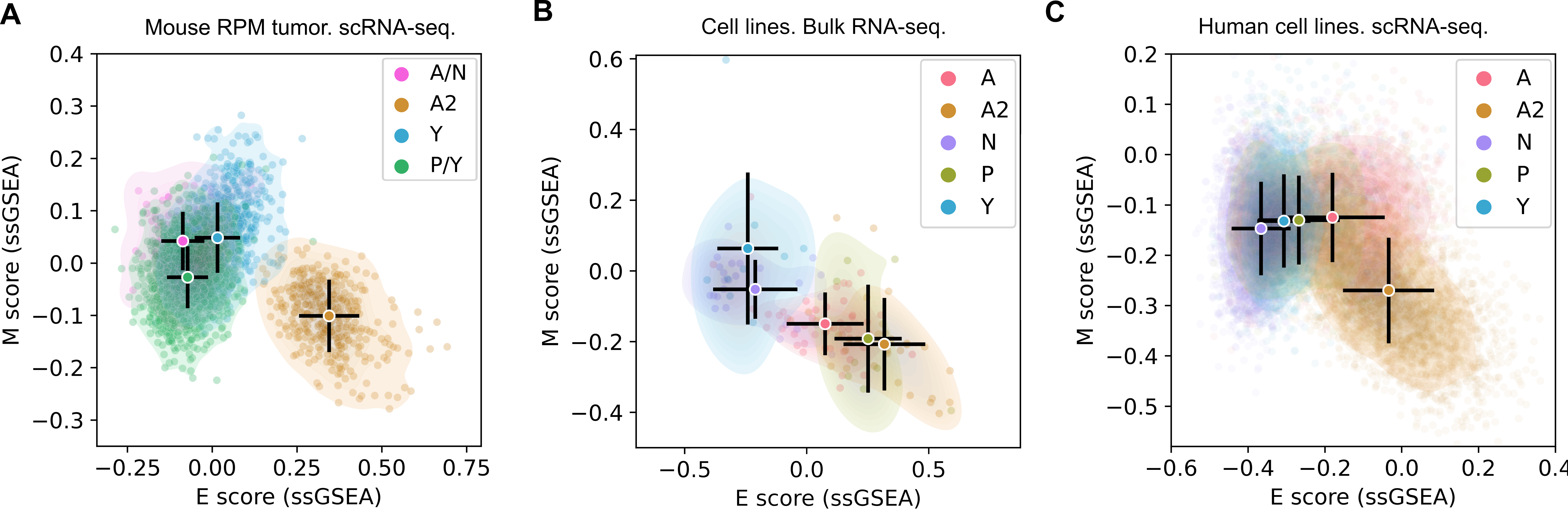

### Figure_S2.png

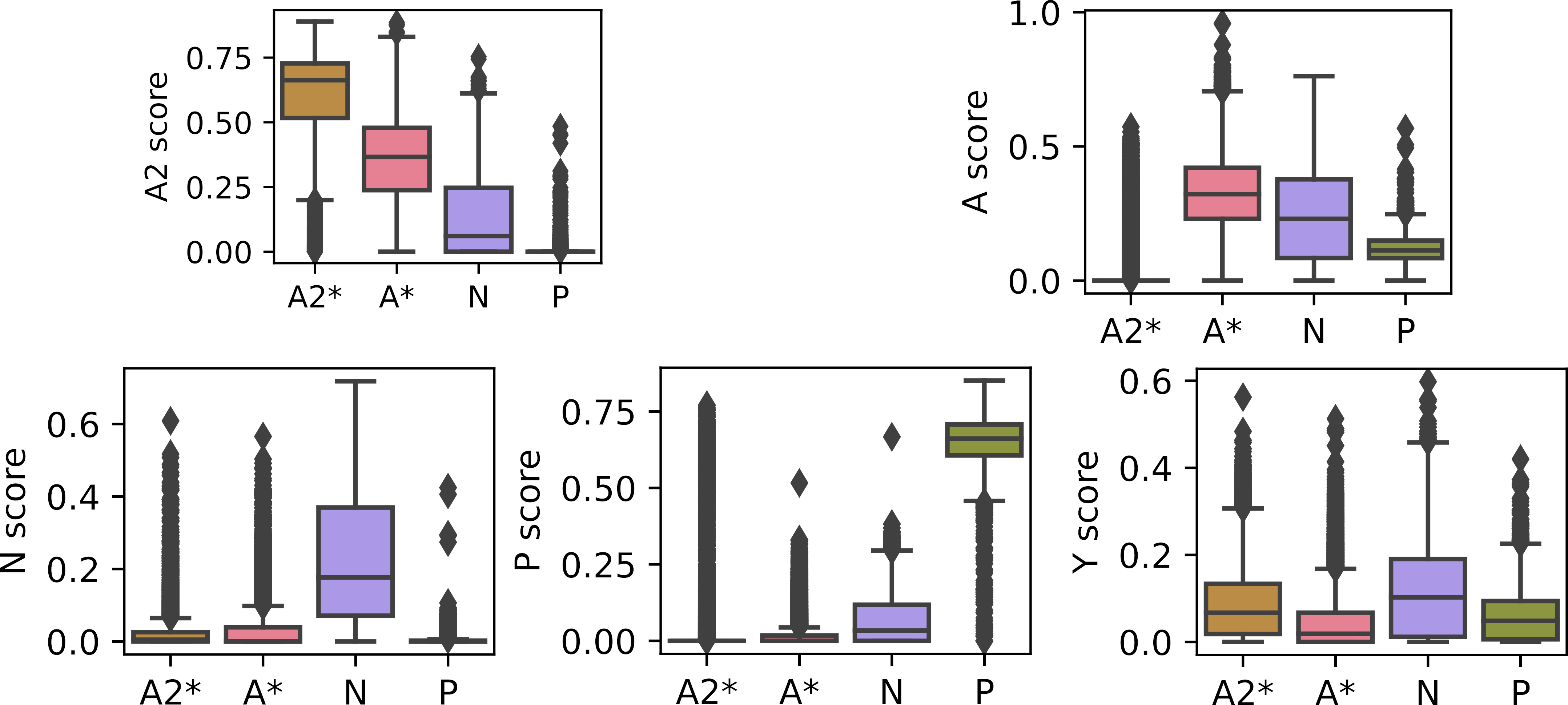

### Figure_S3.png

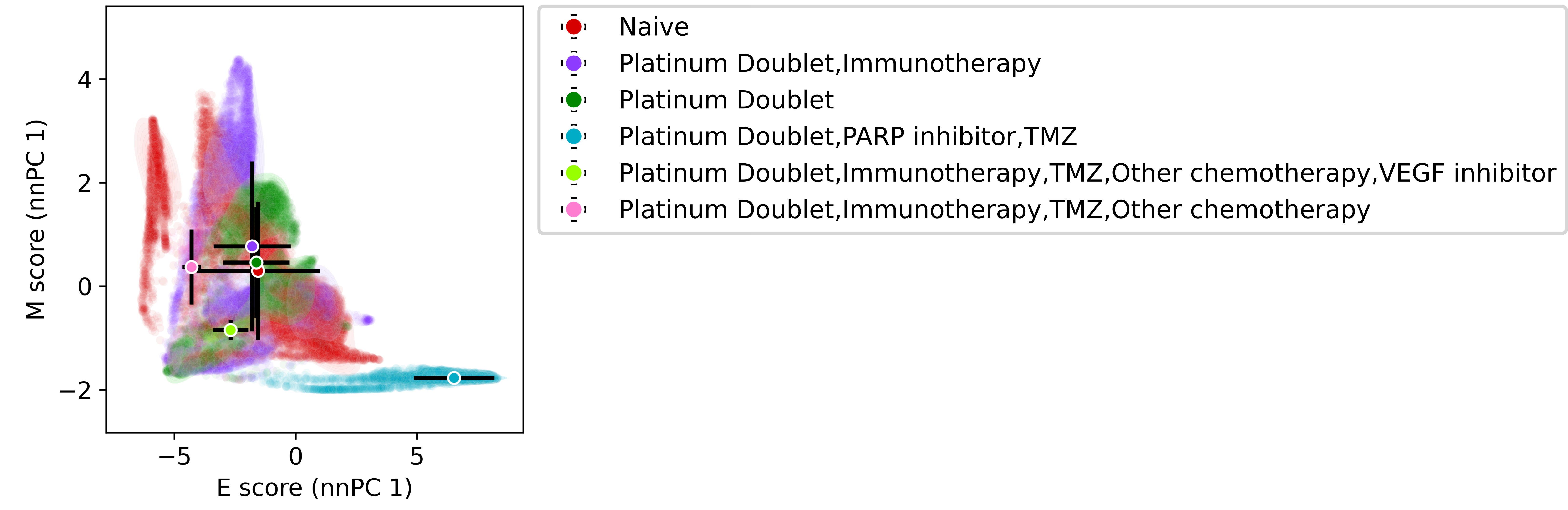

### Figure_S4.png

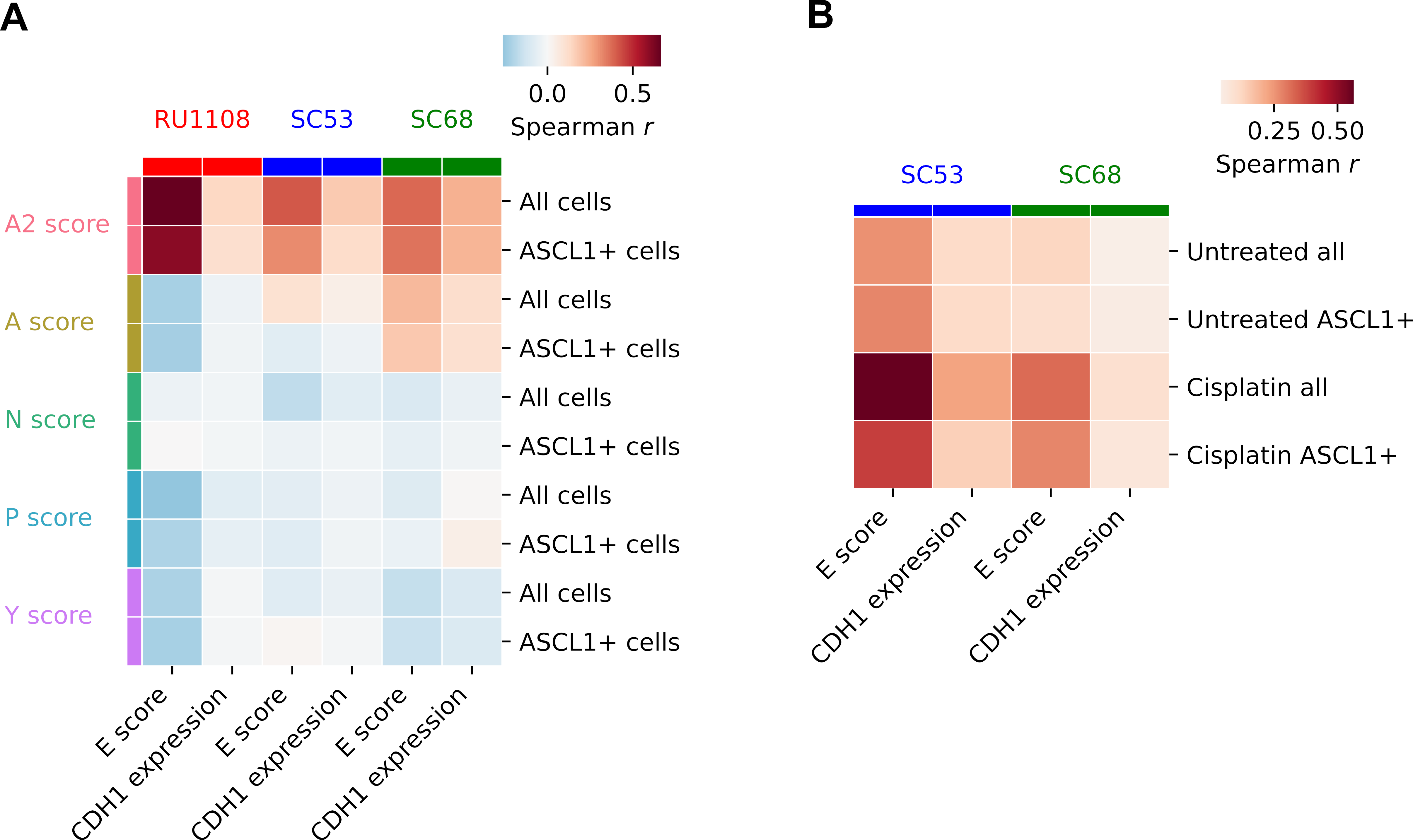
